# Highly pathogenic Avian Influenza A (H5N1) Clade 2.3.4.4b in Wild Birds, Ecuador

**DOI:** 10.1101/2023.10.18.562614

**Authors:** Andrés Carrazco-Montalvo, Lucia Luje, Cristina Rodríguez-Pólit, Antonio Ampuño, Leandro Patiño, Diana Gutiérrez-Pallo, Daniel Alava, Damaris Alarcón-Vallejo, Natali Arguello, Gabriela Echeverría-Garcés, David De La Torre

**Affiliations:** Instituto Nacional de Investigación en Salud Pública “Leopoldo Izquieta Pérez”. Centro de Referencia Nacional de Genómica, Secuenciación y Bioinformática. Quito, Ecuador; Ministerio de Ambiente, Agua y Transición Ecológica. Quito, Ecuador; Instituto Nacional de investigación en Salud Pública “Leopoldo Izquieta Pérez”. Dirección Técnica de Investigación, Desarrollo e Innovación. Guayaquil, Ecuador; Instituto Nacional de investigación en Salud Pública Dr. Leopoldo Izquieta Pérez. Laboratorio Zonal de Influenza y Otros Virus Respiratorios. Quito, Ecuador; Laboratorio de Biología y Genética Molecular - LABIGEN. Quito, Ecuador

**Keywords:** highly pathogenic avian influenza virus, H5N1, clade 2.3.4.4b, wild birds, viruses, zoonoses, influenza, Ecuador

## Abstract

Highly pathogenic avian influenza A(H5N1) virus clade 2.3.4.4b was detected in four wild birds of two species, *Fregata magnificens* and *Sula nebouxii*, on the Ecuadorian Coast. This report highlights the importance of intersectoral collaboration and timely genotyping for monitoring this zoonotic pathogen, especially in regions with a rich biodiversity.

---

Highly pathogenic avian influenza (HPAI) has been recognized for its pandemic potential since the early 2000s (*1*). The avian influenza viruses are categorized based on 16 Hemagglutinin (HA) and nine Neuraminidase (NA) subtypes, and they circulate in natural enzootic cycles primarily in wild aquatic birds. Zoonotic infections and outbreaks of HPAI have been mainly caused by the H5 and H7 subtypes (*2*). From 2003 to July 2023, there have been 878 documented cases of human avian influenza A (H5N1) across 23 countries worldwide (*3, 4*). In January 2023, the World Health Organization (WHO) received the first report of a human infection in Ecuador (*5*).

Since 2021, subtype A (H5N1) clade 2.3.4.4b have caused important outbreaks in poultry and wild birds across Europe, the Middle East, Asia, and North America (*6*). In October 2022, this clade was first reported in South America (*6-7*) causing mass mortality in poultry. Furthermore, recently in 2023, this clade caused illness and mass mortality in wild bird species such as pelicans (*Pelecanus thagus*) and wild mammals like sea lions (*Otaria flavescens*) in Peru and Chile (*8-9*).

To effectively monitor the emergence and spread of new cases among susceptible species and humans, intersectoral collaboration, routine surveillance within the “One Health” context, and genotyping are urgently needed.

## The study

This report is the result of a joint effort within the “One Health” approach involving the Ministry of Environment, Ecuador (MAATE by its Spanish acronym), and the National Institute of Public Health Research “Leopoldo Izquieta Pérez”, Ecuador (INSPI by its Spanish acronym). We report four cases of infection by Influenza A H5N1/HPAI clade 2.3.4.4b causing illness in wild birds from two species: two frigatebirds (*Fregata magnificens*) and two blue-footed boobies (*Sula nebouxii*) sampled in different regions of the Ecuadorian Coast. The samples of frigatebirds (one sick and one dead) were collected on January 2023 from San Vicente-Manabí province, the samples of blue-footed boobies (both sick) were collected on May 2023 from Villamil Playas-Guayas province (Figure 1A-C). Sample collection was carried out by MAATE (Supplementary Material 1).

**Figure 1.**
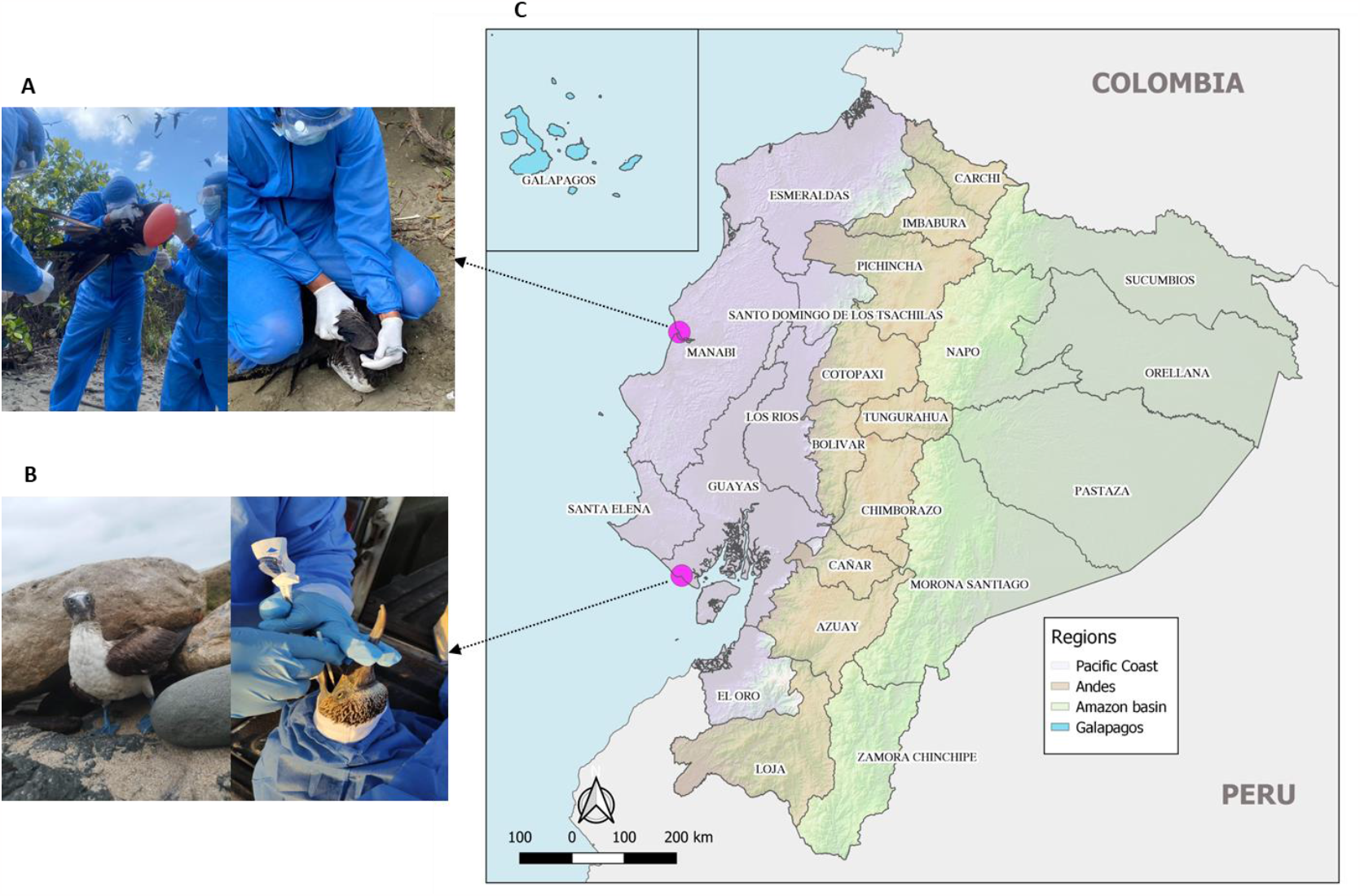
**A)** Frigatebird (*Fregata spp*.) sample collection: tracheal swab. **B)** Blue-footed booby (*Sula nebouxii*) sample collection: tracheal swab. **C)** Map of Ecuador showing collection points: San Vicente-Manabí and Villamil Playas-Guayas.

Tracheal swab samples were diagnosed positive for A/H5N1 using RT-qPCR at a private laboratory in Quito, (LABIGEN). Diagnosis confirmation and genotyping were performed at the INSPI, headquarters in Quito. Samples were confirmed positive for A/H5 at the Zonal Laboratory for Influenza and Other Respiratory Viruses using the gold standard test RT-qPCR. Whole genome sequencing and bioinformatics analysis were conducted at the National Reference Center for Genomics, Sequencing, and Bioinformatics (CRN-GENSBIO by its Spanish acronym). Sequencing was performed on an Illumina MiSeq instrument using the Illumina Respiratory Virus Oligo Panel (RVOP) kit and a standard 150-cycle V2 flow cell. FASTQ files were analyzed using FastQC (*10*) and the adapters were trimmed with Trimmomatic (*11*). Genome assembly was carried out with the reference genome Influenza A/Goose/Guangdong/1/96 (H5N1) (accession number: NC_007362) using the Bowtie2 v2.5.1 tool (*12*). Four whole genome sequences, one from each sample, were obtained and deposited in GISAID (GISAID Accession Numbers: EPI_ISL_17973443, EPI_ISL_17973458, EPI_ISL_18137626, and EPI_ISL_18137671).

The genomes were confirmed to belong to the A H5N1/HPAI clade 2.3.4.4b (Figure 2A). Neuraminidase 1 (N1) was determined using the BLAST (https://www.ncbi.nlm.nih.gov/) and BV-BRC (https://www.bv-brc.org/) tools. Whereas, clade assignment was performed through phylogenetic analysis of 625 sequences of avian influenza viruses, using BV-BRC.

**Figure 2.**
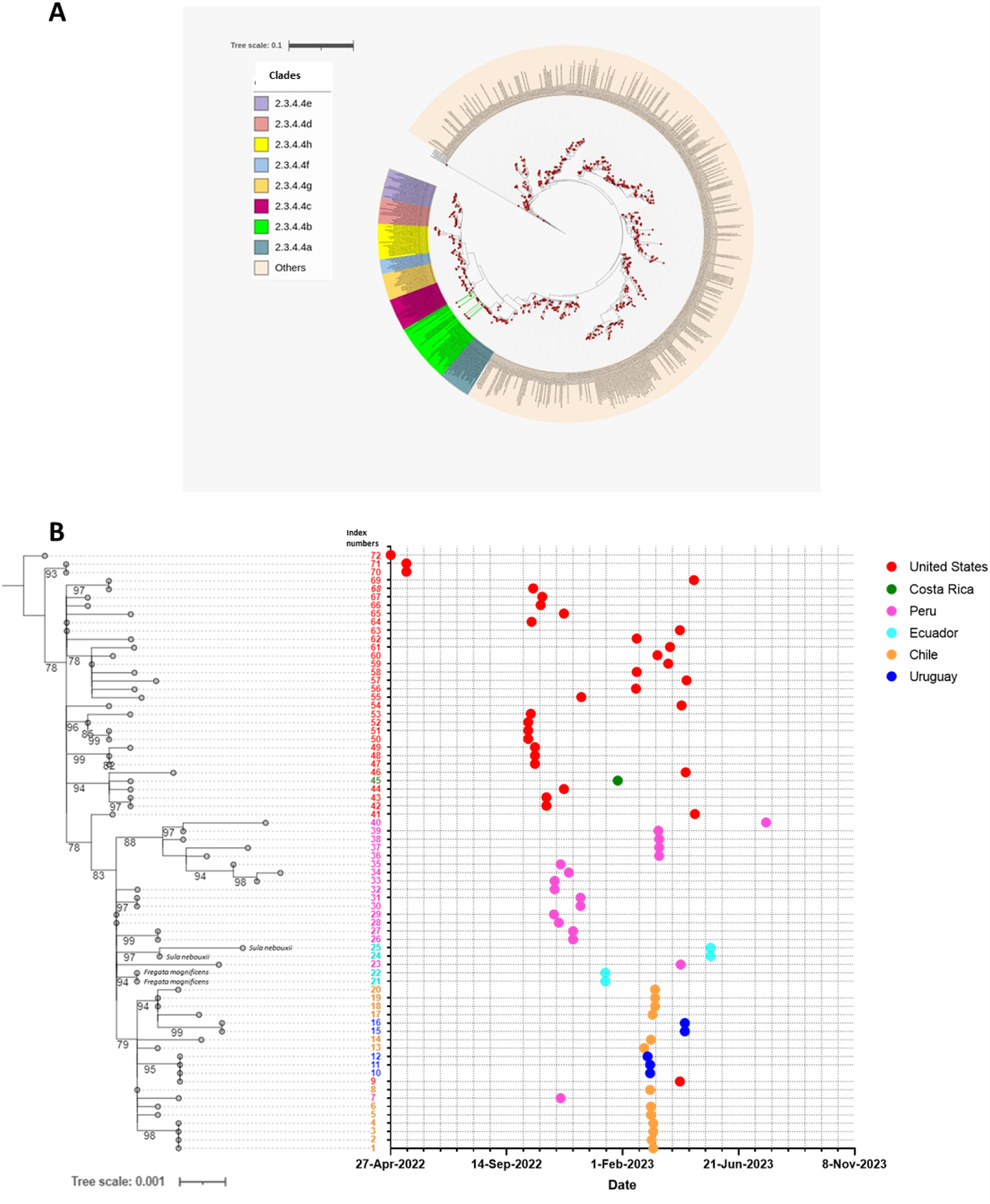
**A)** Clade 2.3.4.4b assignment highlighted in light green. Samples from Ecuador with light green lines. **B)** Phylogenetic tree generated from the analysis of 68 sequences closely related with *Fregata magnificens* and *Sula nebouxii* of Ecuador. On the left, the phylogenetic tree displays Bootstrap values > 70. Samples are labeled with index numbers from 1 to 72 at the end of the branches and is color-coded. These colors correspond to the points on the right, which organizes the samples by country and collection dates.

To identify the most similar sequences, the four obtained genomes from Ecuador were compared with 357,193 sequences available in the EpiFlu database using the AudacityInstant (v0.9) tool (https://gisaid.org/). Out of these genomes, 68 were found to be most closely related. In order to identify the genetic relatedness among the 68 sequences and the genomes of Ecuador, phylogenetic analysis was conducted using the Maximum Likelihood (ML) method in IQ-tree 2.2.0 (http://iqtree.cibiv.univie.ac.at/). GraphPad Prism 8 was used to plot sample collection dates and geographic locations for each sequence (see details for index numbers in Supplementary Material 2). Viral sequences obtained from *Fregata magnificens* (index 21 and 22) and *Sula nebouxii* (index 24 and 25) in Ecuador clustered with viral sequences obtained from birds in Peru (Figure 2B), including Peruvian booby (index 23, collected on April 2023), chickens (index 26, 27, 30, and 31, collected on December 2022), pelicans (index 28 and 29, collected on November 2022), and owl (index 32, collected on November 2022).

Ecuador is a megadiverse country that features a remarkable array of ecosystems, including numerous unique and endemic species within a small area (*13*). There is a critical need for consistent monitoring of avian influenza viruses, especially in the Ecuadorian coast due to the geographical closeness the Galapagos Islands, declared Natural World Heritage by UNESCO. Wild birds can be infected with HPAI without showing signs of illness, and they can carry the disease to new areas when they migrate, potentially exposing domestic animals, poultry and other mammals (*14*). Intersectoral collaboration for wild bird surveillance, HPAI diagnosis, genotyping through whole genome sequencing and correlation with epidemiological data might serve as an important early alert mechanism to control potential outbreaks and could offer insights about the prevalence and progression of these viruses, enabling more proactive responses to reduce the impact on susceptible species (*15*). Genome sequencing is crucial for surveying the emergence and spread of virus subtypes timely. Given the reports of infection in both mammals and humans, there is an urgent need for strengthen preparedness and laboratory networks both locally and regionally to face future epidemics.

## Conclusions

Highly pathogenic avian influenza HPAI A(H5N1) virus clade 2.3.4.4b was identified in tracheal samples of four wild birds in Ecuador: two frigate birds (*Fregata magnificens*) and two blue-footed boobies (*Sula nebouxii*). The rapid spread of this virus in the Americas its ability to cause illness in wild birds and spillover to some mammal species, along with its zoonotic potential, underscores the importance of timely surveillance and prevention. The genomic sequences cited in this study were uploaded to the GISAID database, in order to contribute to the broader effort of early detection, tracking and understanding avian influenza viruses and their different subtypes that may pose a public health risk.

## Acknowledges

Special thanks to the global laboratories for their valuable contributions to open-access sequences to GISAID.

## Ethics Statement

In this study, no avian genetic material was used; exclusively viral genetic material was analyzed. The samples were processed in local laboratories through intersectoral collaboration. Sampling in wild birds involved minimally invasive techniques, restricted to tracheal swabs for epidemiological surveillance.

## Notes

### Competing Interest Statement

The authors have declared no competing interest.

